# A nucleoid-associated protein is involved in the emergence of antibiotic resistance by promoting the frequent exchange of the replicative DNA polymerase in *M. smegmatis*

**DOI:** 10.1101/2023.06.12.544663

**Authors:** Wei L. Ng, E. Hesper Rego

## Abstract

Antibiotic resistance in *M. tuberculosis* exclusively originates from chromosomal mutations, either during normal DNA replication or under stress, when the expression of error-prone DNA polymerases increases to repair damaged DNA. To bypass DNA lesions and catalyze error-prone DNA synthesis, translesion polymerases must be able to access the DNA, temporarily replacing the high-fidelity replicative polymerase. The mechanisms that govern polymerase exchange are not well understood, especially in mycobacteria. Here, using a suite of quantitative fluorescence imaging techniques, we discover that, as in other bacterial species, in *M. smegmatis,* the replicative polymerase, DnaE1, exchanges at a timescale much faster than that of DNA replication. Interestingly, this fast exchange rate depends on an actinobacteria-specific nucleoid-associated protein (NAP), Lsr2. In cells missing *lsr2*, DnaE1 exchanges less frequently, and the chromosome is replicated more faithfully. Additionally, in conditions that damage DNA, cells lacking *lsr2* load the complex needed to bypass DNA lesions less effectively and, consistently, replicate with higher fidelity but exhibit growth defects. Together, our results show that Lsr2 promotes dynamic flexibility of the mycobacterial replisome, which is critical for robust cell growth and lesion repair in conditions that damage DNA.

**Importance:** Unlike many other pathogens, *M. tuberculosis* has limited ability for horizontal gene transfer, a major mechanism for developing antibiotic resistance. Thus, the mechanisms that facilitate chromosomal mutagenesis are of particular importance in mycobacteria. Here, we show that Lsr2, a nucleoid-associated protein, has a novel role in DNA replication and mutagenesis in the model mycobacterium *M. smegmatis*. We find that Lsr2 promotes the fast exchange rate of the replicative DNA polymerase, DnaE1, at the replication fork and is important for the effective loading of the DnaE2-ImuA’-ImuB translesion complex. Without *lsr2*, *M. smegmatis* replicates its chromosome more faithfully and acquires resistance to rifampin at a lower rate, but at the cost of impaired survival to DNA damaging agents. Together, our work establishes Lsr2 as a potential factor in the emergence of mycobacterial antibiotic resistance.

## INTRODUCTION

DNA replication and repair are fundamental to bacterial evolution. The function of DNA replication and repair pathways are key determinants of the balance between genetic stability and mutability. To establish this balance, bacteria encode and regulate numerous DNA polymerases with specialized functions in different conditions (1). Importantly, the emergence of antibiotic resistance in the major human pathogen *M. tuberculosis* is mainly driven by chromosomal mutagenesis, as *M. tuberculosis* has no ability for horizontal gene transfer (2, 3). Thus, the mechanisms regulating polymerase expression and activity are of particular importance for mycobacterial evolution.

In mycobacteria, the fidelity of chromosome replication is established by the major replicative polymerase DnaE1, which possesses endogenous proofreading activity (4). In addition to DnaE1, mycobacteria encode a suite of other polymerases with specialized repair functions that are only expressed under certain conditions, such as oxidative or DNA-damaging stress (5–7). For example, the mycobacterial specialist polymerases DinB1, DinB2, DinB3, and DnaE2 are induced under different conditions and have varying repair activities and mutational signatures (5–10). Of particular importance is the error-prone polymerase DnaE2. It and its accessory proteins ImuA’ and ImuB – collectively referred to as the ‘mutasome’ – are highly expressed in conditions that damage DNA and are the primary drivers of stress-induced mutagenesis *in vitro* and potentially *in vivo* (5, 6, 11). In general, however, the mechanisms that facilitate switching between polymerases at the site of repair are not well understood, especially in mycobacteria (12).

It is known that in non-DNA damaging conditions, in *E. coli*, components within the large multi-protein complex that replicates DNA (the replisome) exchange rapidly, likely when encountering DNA-protein obstacles. More specifically, fluorescence recovery after photobleaching (FRAP) and single molecule studies have revealed that, despite being highly processive *in vitro*, replisome components are highly dynamic within the cell (13). For example, at room temperature, the alpha subunit of Pol III in *E. coli* exchanges with the cytoplasmic pool at a time scale of approximately 4 seconds, suggesting that subunits are constantly being replaced during the 150 minutes it takes to replicate the chromosome at this temperature (13). These data underlie the notion that the replisome is dynamic and flexible, allowing new subunits the opportunity to exchange frequently and suggest that protein obstacles influence the exchange rate of replisome components.

Mycobacteria differ from model bacteria like *E. coli* in several ways. Most notable is their slow growth and long doubling times. Pathogenic mycobacteria, like *M. tuberculosis* and *M. leprae*, are notorious for their long doubling times, which can vary from 18 hours to several weeks. Even so-called fast growers like *M. smegmatis* double on the hour-long timescale compared to the minute-long timescale of commonly studied organisms like *E. coli*. Intriguingly, the duration of chromosomal replication at least partially scales with growth rate: *M. smegmatis* exhibits a longer duration of chromosome replication than *E. coli* (∼100-150 minutes vs ∼41 minutes, respectively) despite the ability of DnaE1 to polymerize DNA at a faster rate than *E. coli* Pol IIIα *in vitro* (4, 14, 15).

These observations led us to wonder if components within the replisome exchange at a different timescale in mycobacteria than in faster-growing bacteria. Using various fluorescence imaging techniques, we find that, as in *E. coli*, the replicative polymerase is exchanged rapidly in *M. smegmatis*. Surprisingly, this fast exchange rate is, in part, mediated by an actinobacteria-specific DNA-binding protein Lsr2, as DnaE1 is exchanged more slowly in cells missing *lsr2*. Interestingly, when the DNA is damaged, ImuB, a key scaffolding protein that recruits the error-prone polymerase into an active complex, is exchanged more rapidly, suggesting ineffective loading of the mutasome. Consistently, loss of *lsr2* results in growth defects in conditions that damage DNA, and reduced mutagenesis. Overall, our data deepen our understanding of DNA metabolism in mycobacteria and provide another unexpected layer to the mechanisms mycobacteria use to mutate and acquire drug resistance.

## RESULTS

### The mycobacterial replisome is highly dynamic

To investigate the dynamics of DnaE1, we created a strain expressing DnaE1-mScarlet from its native promoter. A similar fusion encodes functional DnaE1 (16). To verify that DnaE1-mScarlet reports on the localization of the replisome, we performed time-lapse microscopy and analyzed fluorescence distributions as a function of both cell length and cell cycle (**Fig. 1A, B**). Consistent with the reported placement of the replisome in mycobacteria, DnaE1 localizes at the approximately mid-cell before disappearing, presumably after DNA replication has been completed (16). Towards the end of the cell cycle, DnaE1 reappears at one- and three-quarter positions to initiate replication in newly synthesized sister chromosomes (**Fig. 1A, B**). These data show that DnaE1-mScarlet is reporting on the localization of the replisome in *M. smegmatis*.

**Figure 1.**
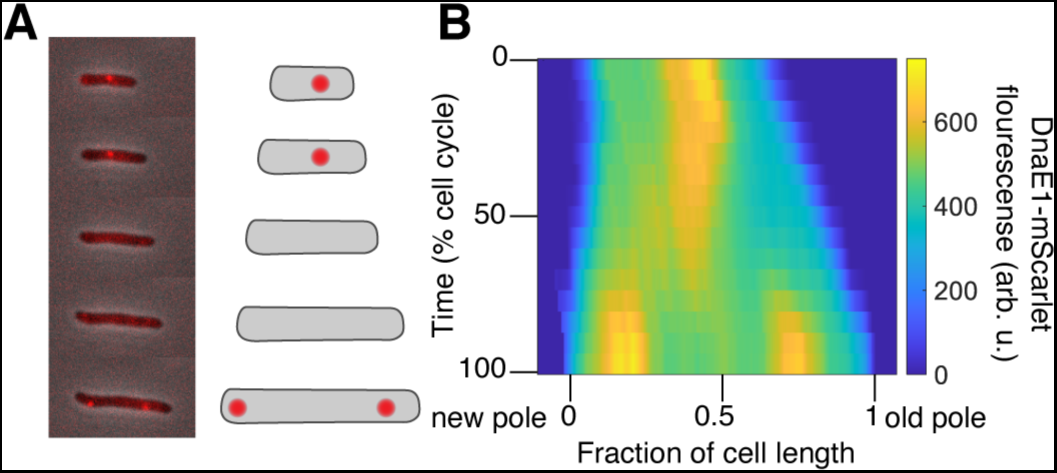
DnaE1 localization over time in *M. smegmatis.* **(A)** A representative cell expressing DnaE1- mScarlet from its native promoter is imaged over time with both phase and fluorescence microscopy. Fluorescent spots appear at mid-cell, disappear, and then re-appear at roughly one- and three-quarters positions at the end of one cell cycle/beginning of another. **(B)** To quantify fluorescence probability distributions over time and space, we constructed kymographs by averaging individual distributions that were normalized to cell cycle time and length (N=43).

To test the exchange rate of DnaE1, we took the same approach as Beattie, TR *et al* and used FRAP to measure the dynamics of recovery of DnaE1 in cells (13). Fast recovery of fluorophores into a bleached spot indicates fast dissociation of bleached molecules. Conversely, slow, or incomplete recovery of non-bleached fluorophores indicates slow dissociation of bleached molecules, and a stably associated complex. To account for differences in the degree of bleaching per cell, we normalized the fluorescence intensity after bleaching to zero, and the final intensity to the maximum recovery we expected based on the percentage of fluorophores bleached. We fit the data to a diffusion exchange reaction and calculated both the timescale of recovery, reflecting the dissociation time of bound subunits, and the maximum recovery, reflecting the mobile fraction of proteins. Imaging was done in the same conditions as above, in which we measured the duration of chromosomal replication to be approximately 100 minutesbased on the time between the appearance and disappearance of DnaE1 (**Fig. 1B, S2B**). We found that, as in *E. coli*, DnaE1 exchanged quickly and nearly completely with a timescale of approximately 8 +/- 3 seconds (mean±SD) (**Fig. 2A-C**). These data are consistent with the model that has emerged in *E. coli,* which suggests that the replisome constantly encounters obstacles that promote the exchange of individual subunits with the cytoplasmic pool.

**Figure 2.**
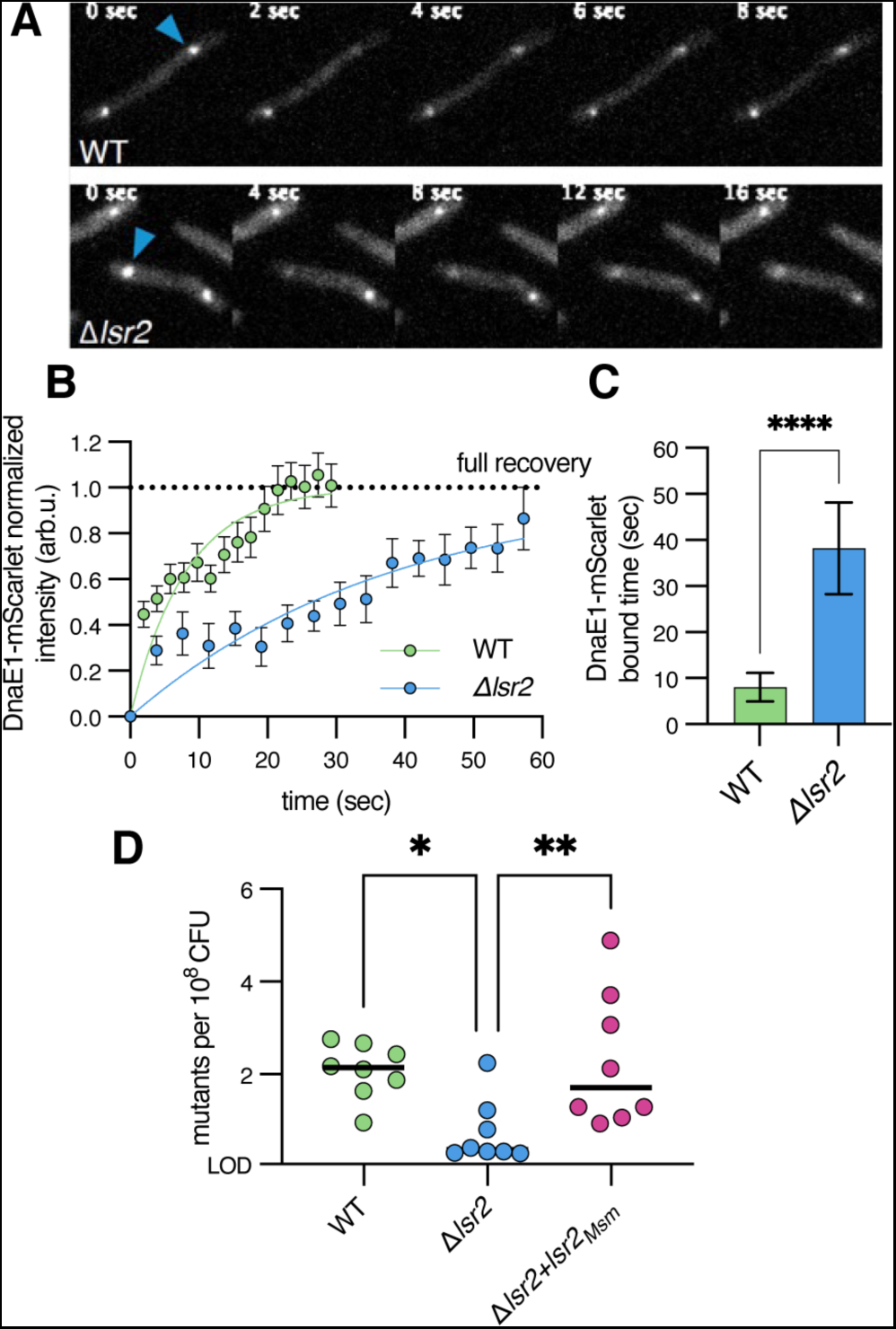
The replicative polymerase is highly dynamic, which is in part mediated by Lsr2. **(A)** Representative time sequence of FRAP of DnaE1-mScarlet in WT (top) and Δ*lsr2* (bottom) imaged at 2 and 4-second intervals, respectively. Blue arrow indicates the location of the focus targeted for bleaching. **(B)** Fluorescence recovery curves for DnaE1-mScarlet (N=21 for WT, 12 for Δ*lsr2*). Solid lines show a one-phase exponential association fit to the data. Individual points show the mean±SEM of fluorescence intensity after bleach correction. Dashed line represents the estimated maximum possible fluorescence recovery. **(C)** DnaE1-mScarlet bound times (mean±SD) obtained from one-phase exponential association fits in panel B for both WT and Δ*lsr*2 for DnaE1. (****), p<0.0001 by unpaired t-test. (**D**) Frequency of spontaneous rifampin-resistant mutants per 10^8^ cfu. LOD refers to the limit of detection. *lsr2* complements were expressed from the native *lsr2* promoter on a phage integrative plasmid. p-values were obtained by one-way ANOVA. (*), p<0.0021 (**), p<0.0002.

### Lsr2 promotes the frequent exchange of DnaE1

In bacteria, nucleoid-associated proteins (NAPS) are important for bacterial chromosome organization and influence replication, transcription, and cell cycle regulation either directly or indirectly. Lsr2 is one of the most well-studied NAPs in mycobacteria. Lsr2 is conserved across mycobacteria and related actinomycetes (17–21). Like H-NS in *E. coli*, Lsr2 primarily binds AT-rich regions of DNA and functions as a global transcriptional repressor (18). Notably, *in vitro* studies show that Lsr2 binding interferes with transcription, inhibits topoisomerase I-dependent supercoil relaxation, and protects DNA from DNase I digestion and H_2_O_2_-mediated degradation (17). Loss of *lsr2* is associated with a wide range of different phenotypes in mycobacteria, including altered cell cycle timing and growth defects upon entering/exiting hypoxic conditions and when exposed to oxidative stress and various antibiotics (17, 22–25). Lsr2 is also involved in resistance against phage infection, protection against reactive oxygen intermediates during macrophage infection, and cells missing *lsr2* show reduced virulence in various hosts, including mice (20, 22, 23, 26).

ChIP-seq data show an abundance of Lsr2 binding sites across the chromosome, and consistent with this, Lsr2 decorates the entire nucleoid in discrete and highly dynamic puncta (24). Interestingly, the major Lsr2 foci are localized near, but not co-localized with, the replisome. We verified this localization pattern in our hands by creating strains that express Lsr2-mScarlet and DnaE1-mGFPmut3. Consistent with prior reports, Lsr2 and DnaE1 exhibit distinct localizations, and although Lsr2 localizes nearby DnaE1 at certain times in the cell cycle, Lsr2 does not co-localize with DnaE1 despite Lsr2’s abundance throughout nucleoid (**Fig. S1**).

Given the importance of Lsr2 to mycobacteria physiology and its localization, we wondered if deletion of *lsr2* would influence replisome exchange dynamics. To that end, we generated an *lsr2* deletion strain expressing DnaE1-mScarlet. Consistent with prior findings using a DnaN reporter, we find that DnaE1 fluorescence disappears from the mid-cell region faster in cells missing *lsr2*, suggesting that DNA replication is faster by approximately 20% (**Fig. S2A, B**) (25). Next, as before, we measured the rate of DnaE1 recovery in *Δlsr2* cells using FRAP (**Fig. 2A**). We observed a much longer recovery time (38 +/- 10 seconds) compared to wild type cells (**Fig. 2B,C**). These data suggest that DnaE1 remains associated with the replisome for longer in cells missing *lsr2*. Alternatively, a longer recovery time in the *lsr2* mutant could be due to reduced DnaE1 access to the replisome (e.g., due to increased access by other polymerases to the replisome). However, reduced access by DnaE1, an essential polymerase in *M. smegmatis*, would likely result in a growth rate difference in the mutant, which we and others do not observe (**Fig. S2C**) (25). Likewise, reduced access by DnaE1 would also increase basal mutation rates, as this polymerase is responsible for faithful chromosome replication in standard laboratory medium (4). Instead, we observe that that Δ*lsr2* cells mutate at decreased frequency as measured by spontaneous resistance to rifampin (**Fig. 2D**), further supporting the notion of increased stability of DnaE1 in Δ*lsr2.* Taken together, these data show that Lsr2 promotes the rapid exchange of DnaE1. In its absence, DnaE1 is more stably associated, and cells replicate their chromosome more quickly and more faithfully.

### Lrs2 is important for the acquisition of mutations by promoting the stable formation of the mutasome

We next considered if Lsr2 also affects polymerase exchange in conditions that elevate the expression of the error-prone polymerase DnaE2 and its associated complex. To test this, we created strains carrying fluorescent proteins fusions to ImuA’, ImuB, and DnaE2 integrated either at a phage site (*imuA’* and *imuB* in Δ*imuAB*) or the chromosomal locus (*dnae2*). When exposed to UV light, these strains acquired resistance to rifampin at a similar rate as wild type, showing that the fusion proteins retain the ability to repair damaged DNA (**Fig. S3**) (27).

ImuB recruits ImuA’ and DnaE2 into an active complex by binding to the β-clamp subunit of the replisome. ImuA’ and DnaE2 bind ImuB but not the β-clamp, and therefore require ImuB to associate with the replisome and catalyze mutations (8, 11, 27, 28). Despite being co-transcribed, ImuA’-mScarlet and mScarlet-ImuB had dramatically different fluorescence intensities: ImuA’-mScarlet was barely detectable above background fluorescence levels, while mScarlet-ImuB was brightly fluorescent (**Fig. S4A,B**). Thus, cellular levels of ImuA’ may be much lower than ImuB, implying that additional post-transcriptional mechanisms exist (e.g. translation regulation or proteolysis) to limit ImuA’ abundance. Alternatively, mScarlet may be cleaved off the ImuA’-mScarlet translational fusion and targeted for degradation. However, consistent with the low abundance of ImuA’, we also observed very low fluorescence of DnaE2-mGFPmut3 (**Fig. S4C**). Others have reported similar results using N-terminal fusions (27). These data suggest that as in *E. coli,* the translesion polymerase abundance may be tightly regulated post-transcriptionally (29). Further, these data suggest that one limiting factor in forming an active mutasome complex may be the low cellular concentrations of ImuA’ and/or DnaE2.

Due to the dim fluorescence of ImuA’ and DnaE2, it was difficult to draw conclusions about their localization or dynamics in cells with or without *lsr2*. Thus, we decided to determine the exchange rate of ImuB. We treated cells with a low concentration of mitomycin (MMC) to induce expression of ImuB and, as before, performed FRAP. In wild type cells, ImuB displayed a much longer timescale of exchange (74 +/- 14 seconds) compared to DnaE1, suggesting ImuB remains associated with the β-clamp for a longer time (**Fig. 3A,B**). The long bound time of ImuB is consistent with the need to recruit low-abundance proteins ImuA’ and/or DnaE2. In cells missing *lsr2*, ImuB exchanged more quickly (46 +/- 6 seconds) than in wild type, at a rate similar to DnaE1-mScarlet (**Fig. 2C**). Thus, as ImuB and DnaE1 both bind the β -clamp, these data suggest that the mutasome complex is less stably associated in cells missing *lsr2*, possibly due to the increased stability of DnaE1.

**Figure 3.**
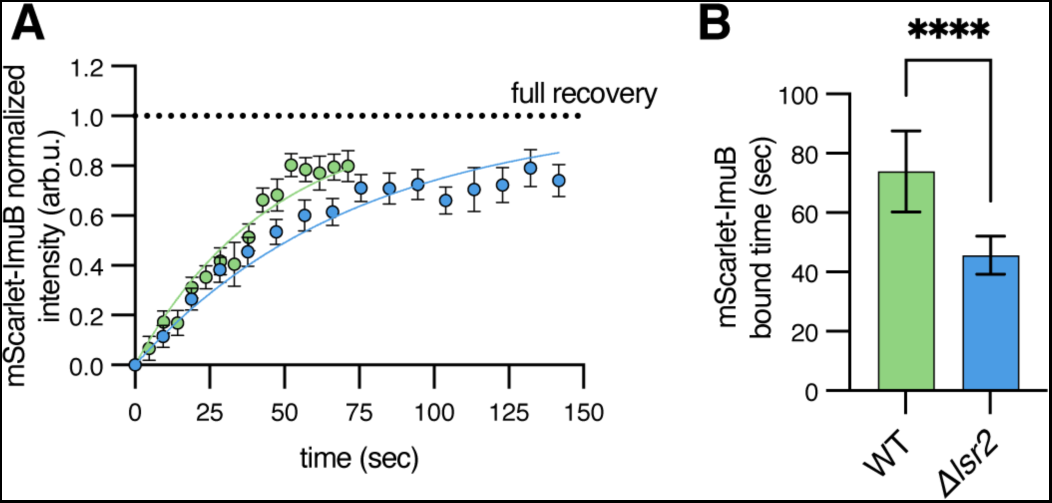
Lsr2 mediates stable loading of ImuB. Cells expressing mScalet-ImuB were treated with 1.56ng/ml mitomycin C for 4 hours before forming FRAP. **(A)** Fluorescence recovery curves for mScarlet-ImuB (N=13 for WT, 10 for Δ*lsr2*). Solid lines show a one-phase exponential association fit to the data. Individual points show the mean±SEM of fluorescence intensity after bleach correction. Dashed line represents the estimated maximum possible fluorescence recovery. **(B)** Bound times (mean±SD) obtained from one-phase exponential association fits from panel A. p<0.0001 by unpaired t-test.

To further test this hypothesis, we reasoned that Δ*lsr2* would phenocopy cells defective in mutasome expression or stability. *M. smegmatis* and *M. tuberculosis* missing ImuA’ and ImuB are more sensitive to MMC treatment and are less likely to mutate when exposed to DNA- damaging agents like ultraviolet light (**Fig. S3A**) (11, 27). Thus, we hypothesized that cells missing *lsr2* would be altered in these key phenotypes. We first confirmed by RNA-seq that the expression of mutasome components or other genes related to the DNA-damage response was unaltered in Δ*lsr2* (**Supplementary Tables 1-9**). Then, to test our model, we grew cells on agar plates with different concentrations of MMC. Deletion of *lsr2* results in a severe growth defect in the presence of MMC, which could be complemented by the expression of *lsr2* from *M. smegmatis* or *M. tuberculosis* (**Fig. 4A, Fig. S5**). Next, we assayed the ability of Δ*lsr2* to acquire resistance to rifampicin when exposed to UV. We chose a dose of UV that resulted in no survival difference between the two genotypes (**Fig. S6**). Consistent with a role in DNA damage response, we found that cells deleted for *lsr2* are approximately 4-5 times less likely to acquire resistance to rifampicin when exposed to UV (**Fig. 4B**). Together, our data show that Lsr2 is involved in the exchange of polymerase at the replication fork in both standard culture medium and in conditions that damage DNA and up-regulate repair pathways.

**FIGURE 4.**
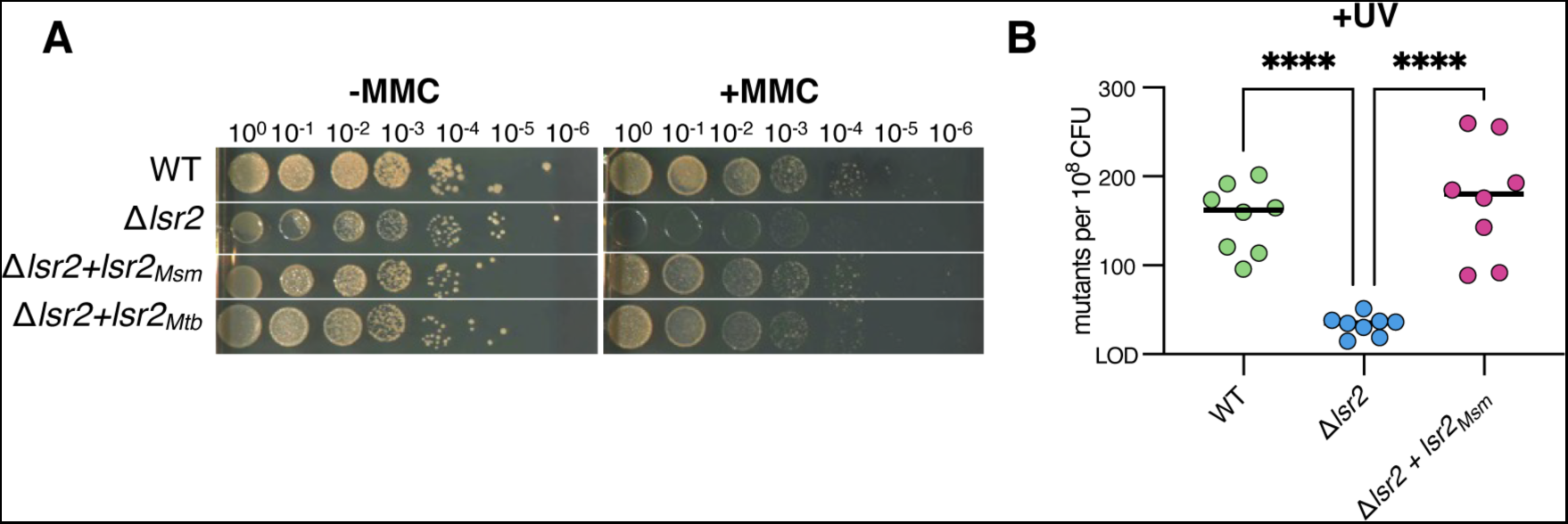
Lsr2 mediates survival to mitomycin and UV-induced mutagenesis. **(A)** Serial dilutions of WT, Δ*lsr2*, and complemented strains on agar plates with or without mitomycin C (1.56 ng/ml). Results shown are representative spots of biological triplicates (**Fig. S5**). **(D)** Frequency of rifampin-resistant mutants per 10^8^ cfu with UV-induced DNA damage (20 mJ/cm^2^). LOD refers to the limit of detection. *lsr2* complements were expressed from the native *lsr2* promoter on a phage integrative plasmid. p-values were obtained by one-way ANOVA (****), p<0.0001.

## DISCUSSION

Lsr2 acts as a transcriptional repressor for many genes on the *M. smegmatis* and *M. tuberculosis* chromosomes (17, 22–24). Additionally, multiple lines of evidence suggest that Lsr2 has other roles aside from its function as a transcription factor (22, 25, 26). Our results implicate Lsr2 as a potential factor in mycobacterial mutagenesis and antibiotic resistance (**Fig. 6**).

**FIGURE 6.**
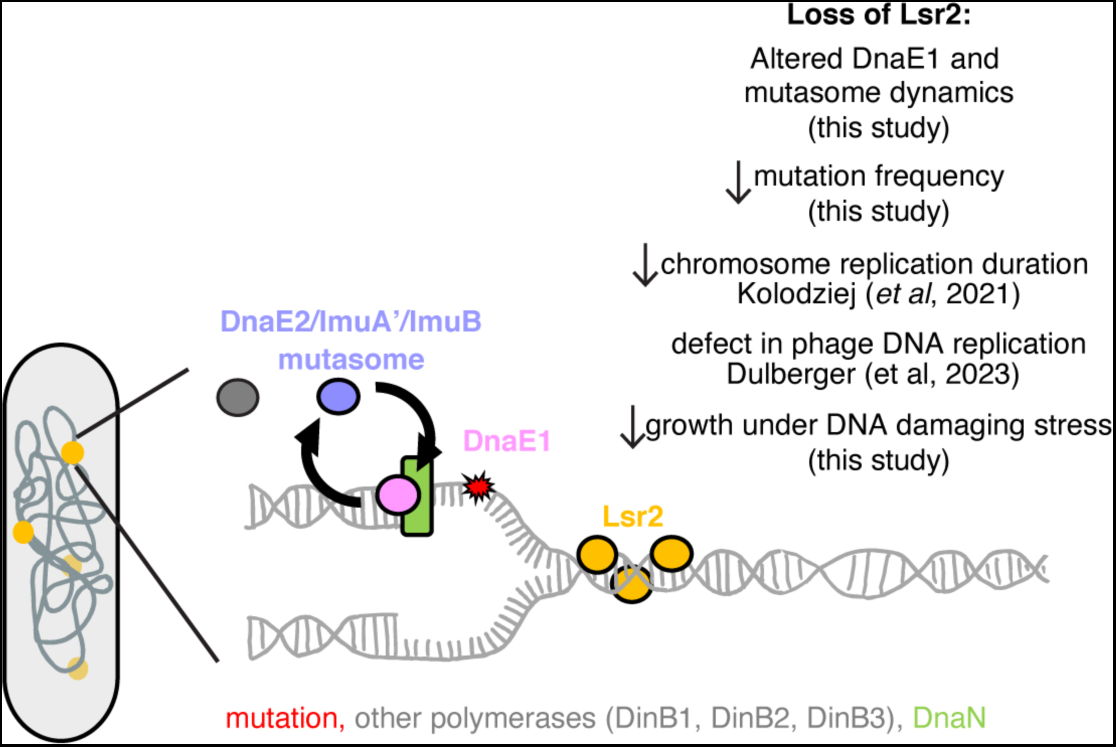
Model of Lsr2 regulation of DNA replication and mutagenesis. Our data suggest that loss of *lsr2* results in a decrease in mutagenesis due to a novel role for Lsr2 in modulating replisome dynamics. Specifically, our data are consistent with a model in which Lsr2 acts as a roadblock on the DNA to promote the exchange of the major replicative polymerase DnaE1 with others in the cytoplasmic pool. In conditions that do not induce DNA damage, these polymerases may include DinB1, which contributes to spontaneous mutagenesis (12). In conditions that damage DNA, the mutasome, which is comprised of DnaE2 and its accessory proteins (ImuA’ and ImuB), is induced and needs to exchange with the replicative polymerase to repair lesions (13, 18, 19). Deletion of Lsr2 reduces the flexibility of the replisome needed for sufficient exchange, thereby resulting in cells that are more reliant on DnaE1, a higher fidelity polymerase that is unable to bypass lesions. Thus, cells missing *lsr2* are impaired in their ability to grow in conditions that damage DNA in their ability to acquire mutations.

While Lsr2 binds to and/or alters the transcription of many genes in both *M. smegmatis* and *M. tuberculosis,* very few phenotypes associated with the deletion mutant have been connected to altered expression of target genes (17, 22, 24). One of the few exceptions is the change in colony morphology, which can be attributed to the increased expression of a putative polyketide synthase (MSMEG_4727) in *M. smegmatis* (24, 30). Why, then, is Lsr2 associated with so many bacterial behaviors, including bacterial growth, the cell cycle, robust infection, survival to hydrogen peroxide and antibiotics, survival in low oxygen, and resistance to phage?

Our results indicate that one of Lsr2’s functions in the cell is to promote the flexibility of the mycobacterial replisome, enabling specialist polymerases like the DnaE2 translesion complex (ImuB/DnaE2/ImuA’) to access the replication fork, leading to efficient, but error-prone, DNA replication and cell growth in conditions that damage DNA. Specifically, our findings support a model whereby in non-DNA damaging conditions, Lsr2 serves as an obstacle that promotes the frequent exchange of DnaE1, allowing other polymerases to exchange into the replisome (**Fig. 6**). This model is consistent with *Δlsr2* having a slightly reduced basal mutation rate, as Lsr2 could be allowing exchange of polymerases besides DnaE1 at a low frequency. Likewise, in DNA damaging conditions, out data suggest a model in which Lsr2-mediated exchange of DnaE1 promotes loading of ImuB and thus DnaE2 and ImuA’. DnaE1 is a more faithful enzyme but is unable to bypass lesions; thus, cells missing *lsr2* grow more slowly under conditions that damage DNA but replicate with higher fidelity.

Numerous studies in eukaryotes and other bacteria show that the rate of replication fork progression is actively regulated, in some cases by machinery that slow down DNA replication, to ensure coordination amongst DNA replication, DNA repair, and transcription (31–34). We hypothesize that Lsr2 may function to regulate replication fork progression in mycobacteria, and that this function may provide a unifying explanation for many of the phenotypes associated with loss of *lsr2,* including slower DNA replication in the host and other environmental conditions likely to assault DNA. Likewise, in conditions that do not significantly damage DNA, like those found in laboratory growth media, loss of *lsr2* leads to faster and higher fidelity chromosomal replication, possibly explaining the emergence of spontaneous *lsr2* mutants in several laboratories (30, 35, 36). Additionally, loss of *lsr2* is associated with increased resistance against infection by certain phages, and fewer and smaller zones of phage DNA replication (26). We speculate that efficient turnover of the bacterial host DNA replication machinery mediated by Lsr2 could be required for robust replication of phage DNA and productive infection.

Intriguingly, Lsr2 is phosphorylated by PknB, a plasma membrane-bound eukaryotic serine/threonine kinase involved in cell growth and division (37). Future research is needed but we speculate that such regulation could be used to coordinate cell wall synthesis and DNA synthesis during environmental changes or even during a typical cell cycle.

Lsr2 may be a promising drug target for preventing the emergence of antibiotic resistance in mycobacteria. Our work adds to the growing body of evidence Lsr2 is at the nexus of several evolutionary tradeoffs, including fast chromosome replication, mutagenesis, replisome flexibility, and phage resistance. Indeed, loss of Lsr2 results in severe growth defects in *M. tuberculosis* and *M. abscessus in vivo* (20, 23). Thus, we speculate that targeting Lsr2 therapeutically may help treat drug-susceptible infections, while preventing the emergence of drug resistance.

## MATERIALS AND METHODS

### Strains and growth conditions

All mycobacterial strains are derivatives of *Mycobacterium smegmatis* MC2 155. Cells were grown in Middlebrook 7H9 broth supplemented with 0.05% (v/v) Tween80, ADC, and 0.2% (v/v) glycerol for liquid culture. LB agar was used as solid media. Liquid cultures were grown at 37 °C on a rotator. Agar plates were incubated at 37 °C. If applicable, antibiotics were added at the following final concentrations: hygromycin B (50 ug/mL), kanamycin (25ug/mL), nourseothricin (20 ug/ml), zeocin (20 ug/mL), and gentamicin (5 ug/ml).

Plasmid DNA for transformation into *M. smegmatis* was prepared from *E. coli* (DH5a, XL1). If applicable, antibiotics were added at the following final concentrations: hygromycin B (100 ug/mL), kanamycin (50 ug/mL), and nourseothricin (40 ug/ml). Plasmid constructs were generated using Gibson cloning and phage-based integrative vectors. Fluorescent protein fusions were generated using mGFPmut3 and mScarlet-I variants.

Genetic modifications to the native locus were generated as previously described using the mycobacterial Che9c phage RecET recombination system (38). To construct a knockout, we designed a dsDNA substrate encoding a zeocin resistance cassette flanked by loxP sites and ∼500bp of homology on either end of the locus of interest. The dsDNA substrate was then electroporated into *M. smegmatis* strains carrying a plasmid encoding isovaleronitrile-inducible expression of RecET. Colonies were selected by zeocin resistance and screened by PCR using primers annealing to regions flanking dsDNA insertion. In some cases, the zeocin resistance cassette was removed by Cre-Lox recombination. Strains and primers are listed in Tables S12 and S13, respectively.

### Fluorescence microscopy

All microscopy experiments were performed on a Nikon TI-E inverted wide-field microscope equipped with the Nikon Perfect Focus System, an ORCA-Flash4.0 Digital CMOS Camera (Hamamatsu) and an environmental control chamber maintained at 37 °C.

#### Time-lapse and kymograph analysis

Time-lapse imaging was done using a 60X Oil 1.45NA Plan Apochromat phase-contrast objective. Bacterial samples were loaded and grown in a B04A microfluidic plate (CellAsic ONIX, B04A-03-5PK). Phase and fluorescence images were acquired every 30 minutes using NIS Elements software. Fluorescence images were taken sequentially using a 470 nm excitation laser and 515/30 emission filter for msfGFP and mGFPmut3, and a 555 nm excitation laser and a 595/40 emission filter for mScarletI.

Kymograph analysis was performed using a custom pipeline for segmenting and tracking cells, and extracting fluorescence profiles. Cell segmentation was performed on the phase image using a U-Net network trained for segmenting mycobacterial cells (39). Semi-automated cell tracking and extraction of fluorescence profiles from background-subtracted images was done using a custom MATLAB program and open-source image analysis software Fiji (40). Average kymographs were generated by 2D interpolation of at least twenty individual kymographs.

#### FRAP

FRAP experiments were done using a 60X oil DIC objective. Bacterial cells were imaged under 7H9 agar pads (1% agar, 0.2% (v/v) glycerol) in 50 mm glass-bottomed dishes.

Fluorescent spots were manually selected for localized bleaching with a focused 405-nm laser. Two pre-bleach fluorescence images were acquired: one to identify and center a target focus, and a second for quantification of total pre-bleach fluorescence. Fluorescence recovery was monitored by acquisition at pre-defined intervals (2 to 10 sec) following photobleaching using an exposure time of 200 ms.

Cell outlines for determination of total pre-bleach fluorescence were manually drawn using the brightfield image. Fluorescence images were background subtracted and corrected for photobleaching. To quantify the fluorescence of bleached foci over time, a circular ROI was manually drawn around the bleached foci and intensity profiles over time were obtained using Fiji (40). Fluorescence recovery curves were generated by first normalizing fluorescence intensity profiles to pre-bleach focus fluorescence. The post-bleach value was then subtracted from all time points. Maximum recovery was estimated by calculating the total amount of fluorescence in the cell after photobleaching outside the focus. Finally, all time points were divided by the estimated maximum recovery value, such that a value of 0 and 1 correspond to no recovery and maximum recovery, respectively.

### RNA-seq of WT *M. smegmatis* and Δ*lsr2*

WT *M. smegmatis* and Δ*lsr2* were grown in triplicate at 37°C to OD600 ∼0.7-1.0. For mitomycin treatment, 1 ml of cells were added to 4 ml of 7H9 containing 100 ng/ml mitomycin C for a final concentration of 80 ng/ml mitomycin C. Approximately 2-4 ml of cells were harvested at each timepoint and incubated with 2 volumes of RNAprotect (Qiagen) at room temperature for 5 minutes prior to centrifugation for 5 minutes at 4000xg 4°C. The supernatant was discarded, and the cells were then immediately used for RNA extraction or stored at −80°C.

RNA extraction was performed using the Qiagen RNeasy Kit (74104). Cells were resuspended in 500 ul of RLT buffer (Qiagen) with 2-mercaptoethanol and homogenized by bead beating for 3 minute pulses at maximum speed, three times, with 5 minutes of rest on ice in-between. The lysate was then centrifuged at maximum speed for 1 minute, and the supernatant was transferred to a new tube. The remainder of the RNA extraction was performed using the Qiagen RNeasy Kit according to manufacturer instructions. RNA quality was verified using a bioanalyzer (Agilent Technologies TapeStation 2200). DNase treatment and ribosomal RNA depletion were performed at the Microbial Genome Sequencing Center before library preparation and sequencing.

### Mutation frequency analysis

10 ml of bacterial cells were grown to OD600 ∼0.7-1.0. 5 ml of cells were UV irradiated (20 mJ/cm2 UV; Stratalinker UV crosslinker), while the remaining 5 ml of cells were used for the untreated control. Treated and untreated samples were then added to an equal volume of fresh 7H9 media and outgrown for 3 hours at 37°C. 100 ul liquid culture was used for CFU determination, and the remainder of the culture was plated onto Rif-containing LB agar plates (200 ug/ml) for mutant detection. All mutation frequency measurements were performed using 8 biological replicates. Rif-resistant colonies were enumerated after 5-7 days of incubation at 37°C. CFU plates were read after about 2-3 days of growth on non-selective media. Mutation frequencies were calculated by the number of Rif-resistant colonies per ml of culture by the CFU/ml.

### Spot assay

Bacterial cells were grown to OD600 ∼0.7-1.0, normalized to the same OD, and then 10-fold serially diluted. 5 ul of culture were spotted on LB agar containing varying concentrations of antibiotic. Plates were imaged after incubation at 37°C for 2-3 days. All assays were done in biological triplicate.

## RESOURCE AVAILABILITY

### Lead contact

Further information and requests for resources and reagents should be directed to and will be fulfilled by the lead contact, Dr. Hesper Rego.

### Materials availability

All unique/stable reagents generated in this study are available from the lead contact.

### Data availability

RNA-seq sequencing data are deposited to the Short Read Archive (SRA) under project number: PRJNA1060683

## ACKNOWLEDGMENTS

We thank members of the Rego lab for helpful suggestions, and Dr. Eduardo Groisman for discussions. This work was supported by NIH R01 AI148255 and the Searle and Pew Foundations to EHR.

## AUTHOR CONTRIBUTIONS

WLN and EHR conceived experiments and analyzed data. WLN performed experiments. WLN and EHR wrote the manuscript. EHR secured funding for the work.

---

**TABLES S1-6.** All quantified RNA-seq read counts for WT and Δ*lsr2*, in triplicate, treated with mitomycin-C at 0, 1, 3, 6, or 24 hours.

*see attached

**TABLES S7-11.** Differentially expressed genes in WT and Δ*lsr2* treated with mitomycin-C 0, 1, 3, 6, or 24 hours

*see attached

**TABLE S12.**
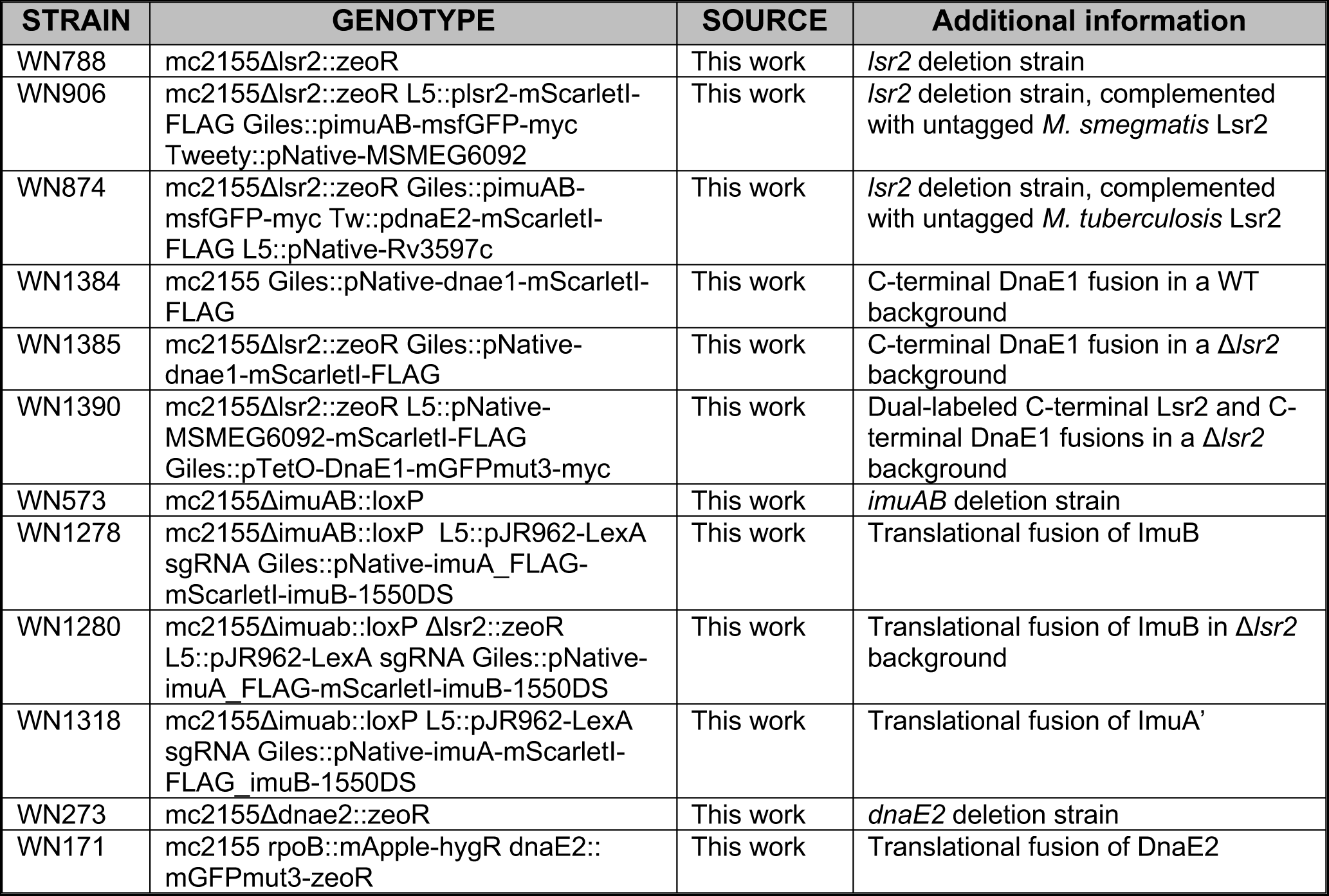
Strain list.

| STRAIN | GENOTYPE | SOURCE | Additional information |
| --- | --- | --- | --- |
| WN788 | mc2155Δl<sub>sr</sub>2::zeoR | This work | *l<sub>sr</sub>2* deletion strain |
| WN906 | mc2155Δl<sub>sr</sub>2::zeoR L5::p<sub>l<sub>sr</sub>2</sub>-mScarletl-FLAG Giles::p<sub>imuAB</sub>-msfGFP-myc Tweety::pNative-MSMEG6092 | This work | *l<sub>sr</sub>2* deletion strain, complemented with untagged *M. smegmatis* L<sub>sr</sub>2 |
| WN874 | mc2155Δl<sub>sr</sub>2::zeoR Giles::p<sub>imuAB</sub>-msfGFP-myc Tw::p<sub>dnaE2</sub>-mScarletl-FLAG L5::pNative-Rv3597c | This work | *l<sub>sr</sub>2* deletion strain, complemented with untagged *M. tuberculosis* L<sub>sr</sub>2 |
| WN1384 | mc2155 Giles::pNative-dnae1-mScarletl-FLAG | This work | C-terminal DnaE1 fusion in a WT background |
| WN1385 | mc2155Δl<sub>sr</sub>2::zeoR Giles::pNative-dnae1-mScarletl-FLAG | This work | C-terminal DnaE1 fusion in a Δ*l<sub>sr</sub>2* background |
| WN1390 | mc2155Δl<sub>sr</sub>2::zeoR L5::pNative-MSMEG6092-mScarletl-FLAG Giles::pTetO-DnaE1-mGFPmut3-myc | This work | Dual-labeled C-terminal L<sub>sr</sub>2 and C-terminal DnaE1 fusions in a Δ*l<sub>sr</sub>2* background |
| WN573 | mc2155ΔimuAB::loxP | This work | *imuAB* deletion strain |
| WN1278 | mc2155ΔimuAB::loxP L5::pJR962-LexA sgRNA Giles::pNative-imuA_FLAG-mScarletl-imuB-1550DS | This work | Translational fusion of ImuB |
| WN1280 | mc2155Δimuab::loxP Δl<sub>sr</sub>2::zeoR L5::pJR962-LexA sgRNA Giles::pNative-imuA_FLAG-mScarletl-imuB-1550DS | This work | Translational fusion of ImuB in Δ*l<sub>sr</sub>2* background |
| WN1318 | mc2155Δimuab::loxP L5::pJR962-LexA sgRNA Giles::pNative-imuA-mScarletl-FLAG_imuB-1550DS | This work | Translational fusion of ImuA' |
| WN273 | mc2155ΔdnaE2::zeoR | This work | *dnaE2* deletion strain |
| WN171 | mc2155 rpoB::mApple-hygR dnaE2::mGFPmut3-zeoR | This work | Translational fusion of DnaE2 |

**TABLE S11.** Primer list

*see attached

**Figure S1.**
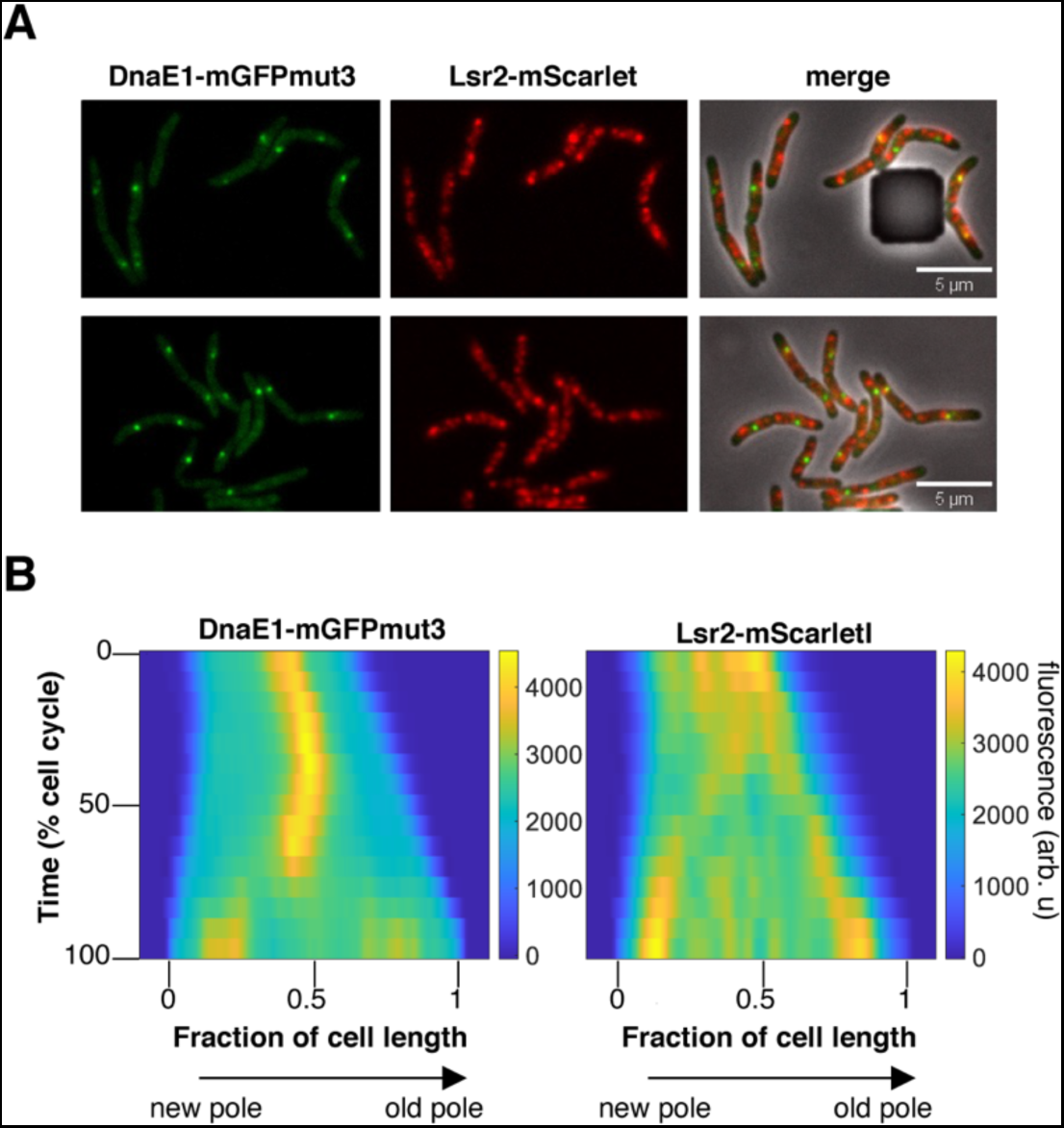
DnaE1 and Lsr2 do not co-localize. **(A)** Microscopy images of cells expressing DnaE1-mGFPmut3 and Lsr2-mScarlet in Δ*lsr2*. **(B)** Average kymographs for each fusion (N=28 cells).

**Figure S2.**
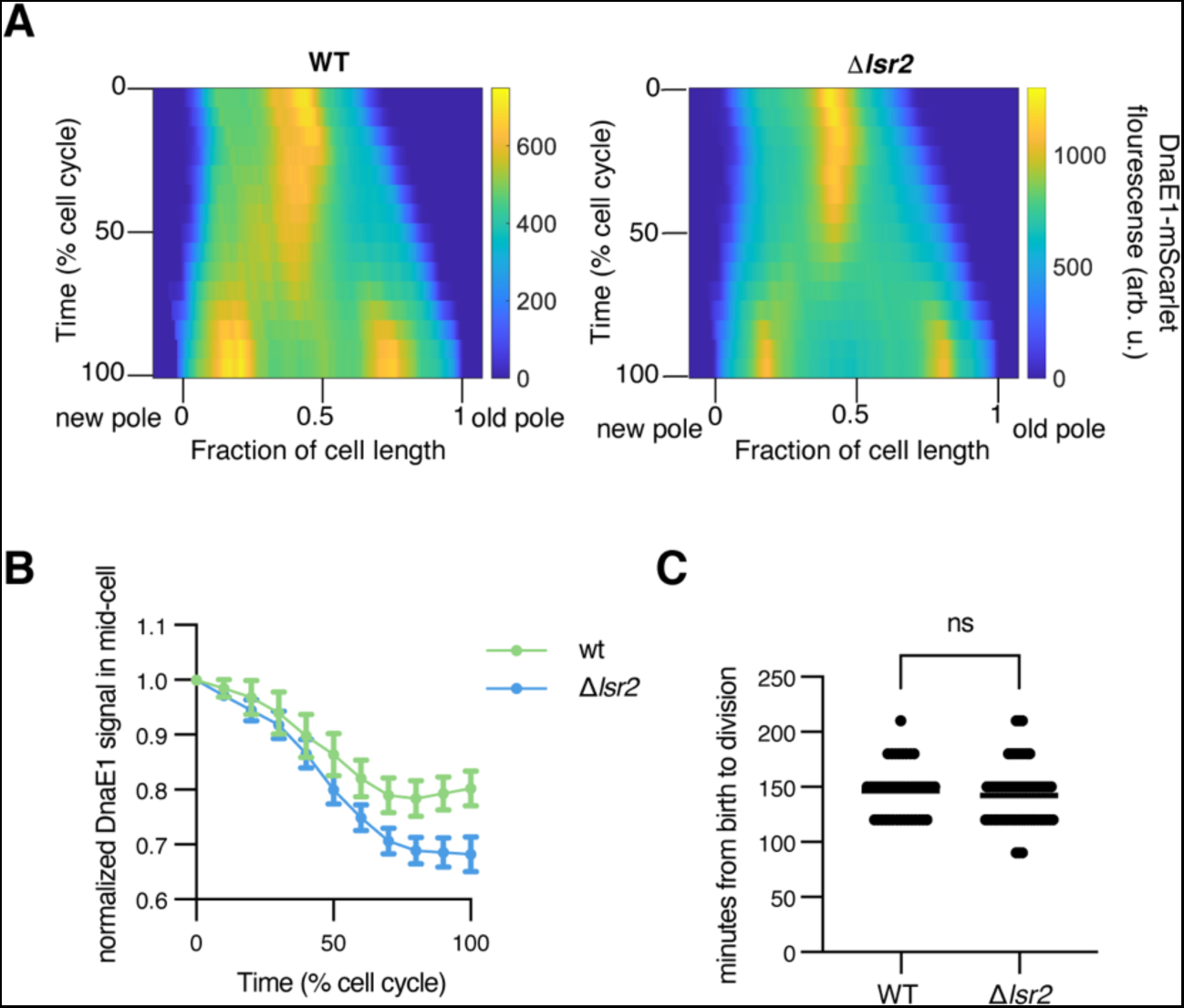
**(A)** Kymograph of DnaE1-mScarlet expressed from the native promoter on a phage integrative plasmid in WT (left, N=43) or Δ*lsr2* (right, N=59). WT is the same as main text Figure 1B for comparison. **(B)** From the cells shown in panel A, the fluorescence at mid-cell from was normalized to time 0 and plotted over time. **(C)** Time between birth and division as determine by visible invaginations on the phase contrast images was measured for both WT and Δ*lsr2* cells.

**Figure S3.**
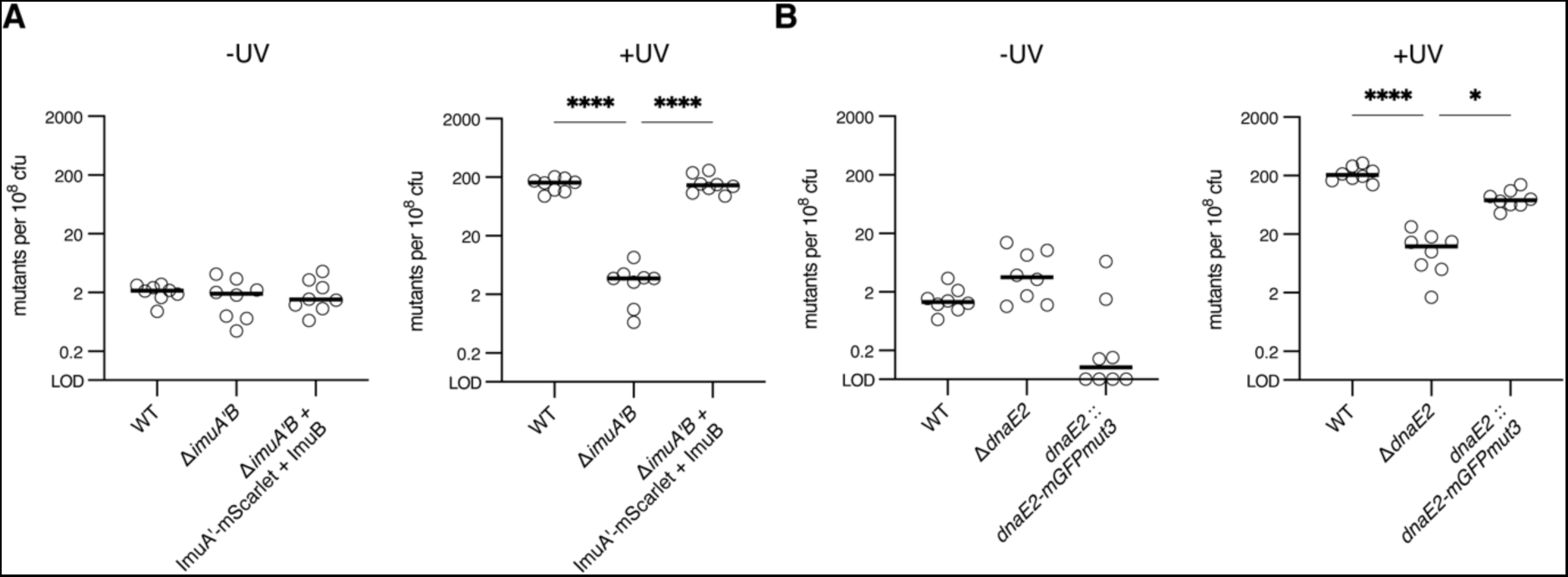
ImuA’ and DnaE2 translational fusions maintain functionality in UV-induced mutagenesis. Frequency of rifampin-resistant mutants per 10^8^ cfu without or with UV-induced DNA damage (20 mJ/cm^2^ UV). **(A)** Results shown are for WT, *imuA’B* deletion, or *imuA’B* deletion complemented with a fluorescent translational fusion of ImuA’ and ImuB. **(B)** Results shown are for WT, *dnaE2* deletion, and *dnaE2* deletion complemented with a fluorescent translational fusion of DnaE2. Complements are expressed from the native promoter on a phage integrative plasmid. LOD refers to the limit of detection. p-values were obtained by one-way ANOVA. 0.1234 (ns), 0.0332 (*), 0.0021 (**), 0.0002 (***), <0.0001 (****).

**Figure S4.**
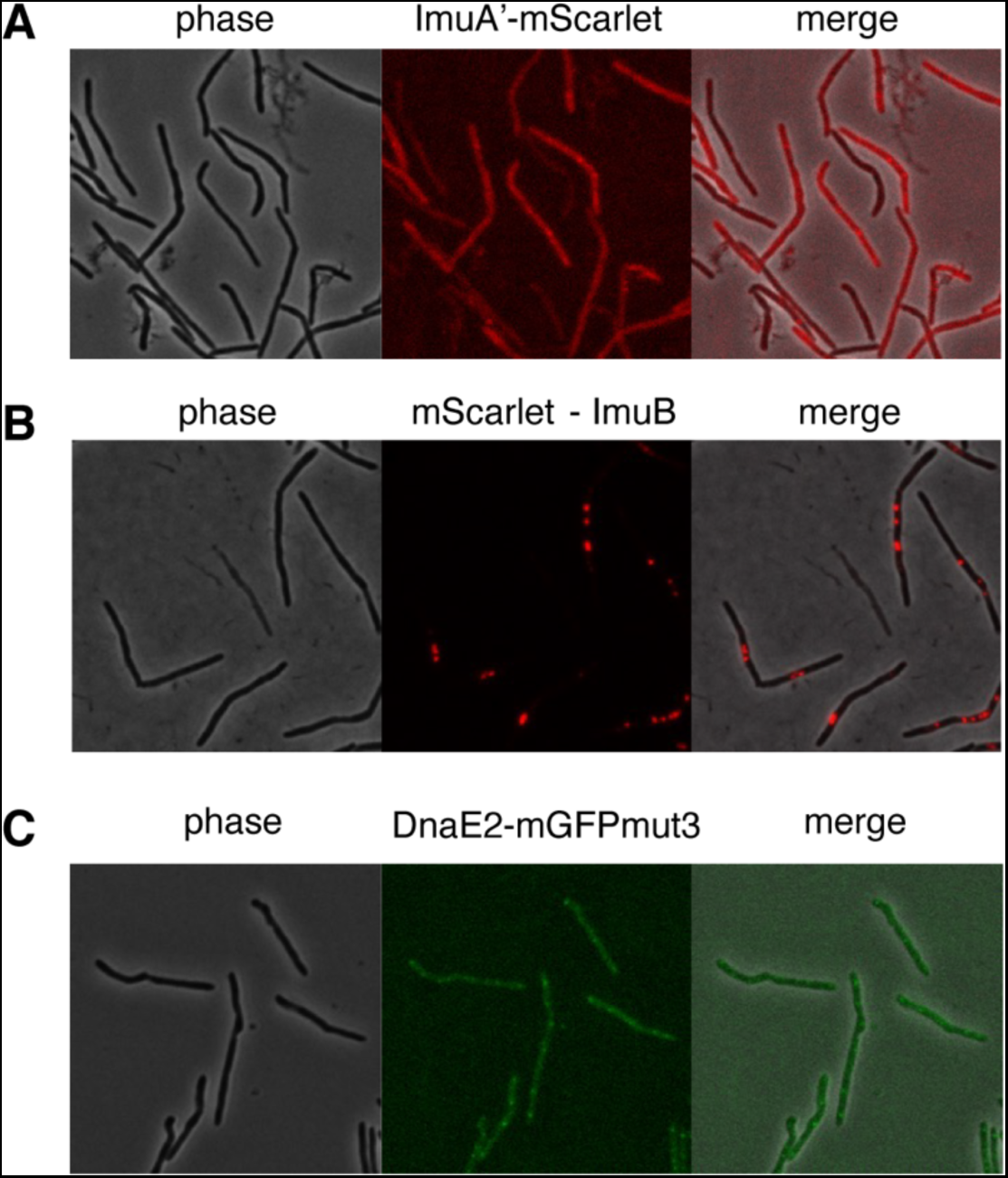
Microscopy images of mutasome translational fusions. **Images shown are of (A)** *imuA’B* deletion mutant complemented with a fluorescent translational fusion of ImuA’ and ImuB, **(B)** *imuA’B* deletion mutant complemented with ImuA and a fluorescent translational fusion of ImuB, and **(C)** *dnaE2* deletion mutant complemented with a fluorescent translational fusion of DnaE2. Complements are expressed from the native promoter on a phage integrative plasmid. DNA damage was induced by mitomycin C treatment (80 ng/ml) for 4-6 hours. Images were acquired using 1 second **(A, B)** or 500 ms **(C)** exposure times.

**Figure S5.**
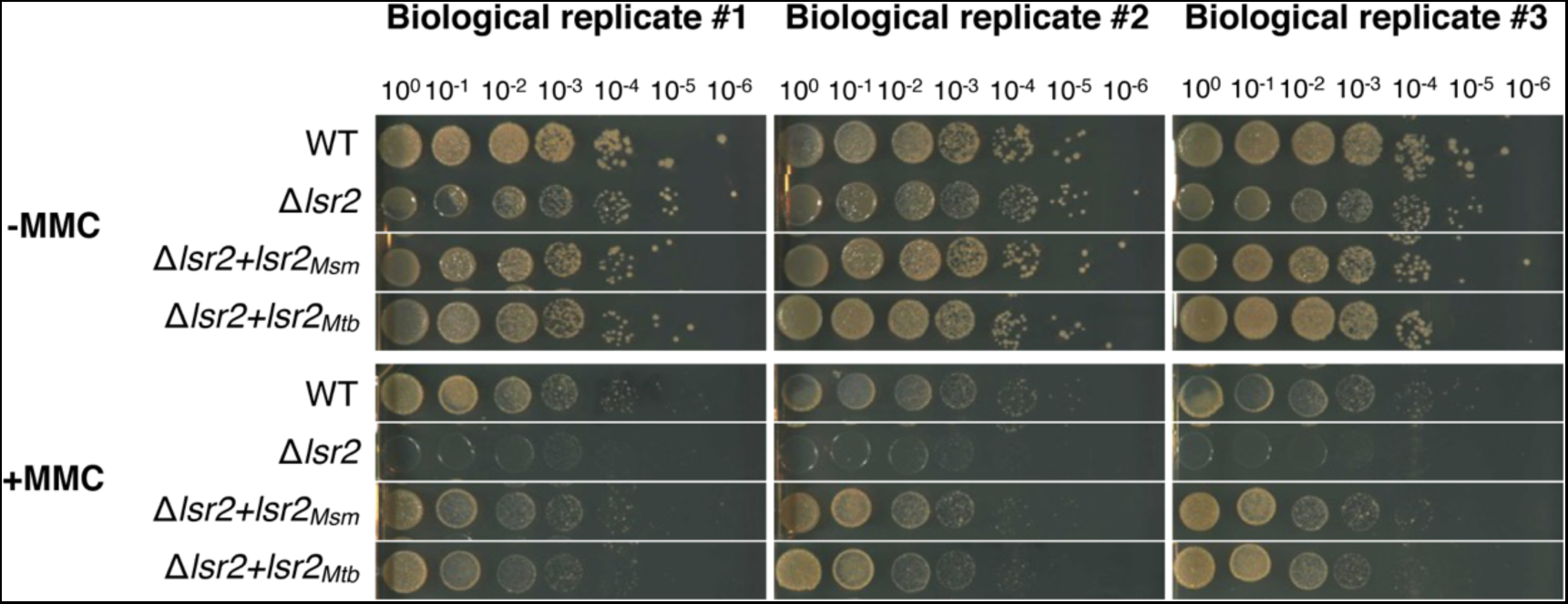
Loss of *lsr2* leads to growth defects on mitomycin C. Serial dilutions of WT, Δ*lsr2*, and complemented strains on agar plates with or without mitomycin C (1.56 ng/ml) in biological triplicate. *M. smegmatis* and *M. tuberculosis lsr2* complements were expressed from the native *lsr2* promoter on a phage integrative plasmid.

**Figure S6.**
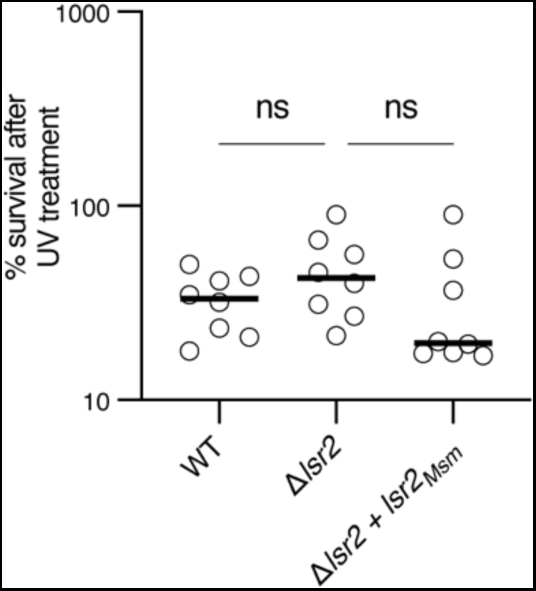
Loss of *lsr2* is not associated with a decrease in survival following UV treatment. WT, Δ*lsr2*, and complement, were treated with 20 mJ/cm^2^ UV and plated for cfu. Percent survival is the ratio of cfu/ml for UV-treated cells divided by the cfu/ml for untreated cells multiplied by 100. p-values were obtained by one-way ANOVA. 0.1234 (ns), 0.0332 (*), 0.0021 (**), 0.0002 (***), <0.0001 (****).

